## Supplemental figures S1, S2 for "Predictive Immunoinformatics Reveal Promising Safety and Anti-Onchocerciasis Protective Immune Response Profiles to Vaccine Candidates (Ov-RAL-2 and Ov-103) in Anticipation of Phase I Clinical Trials"

### Slide 1
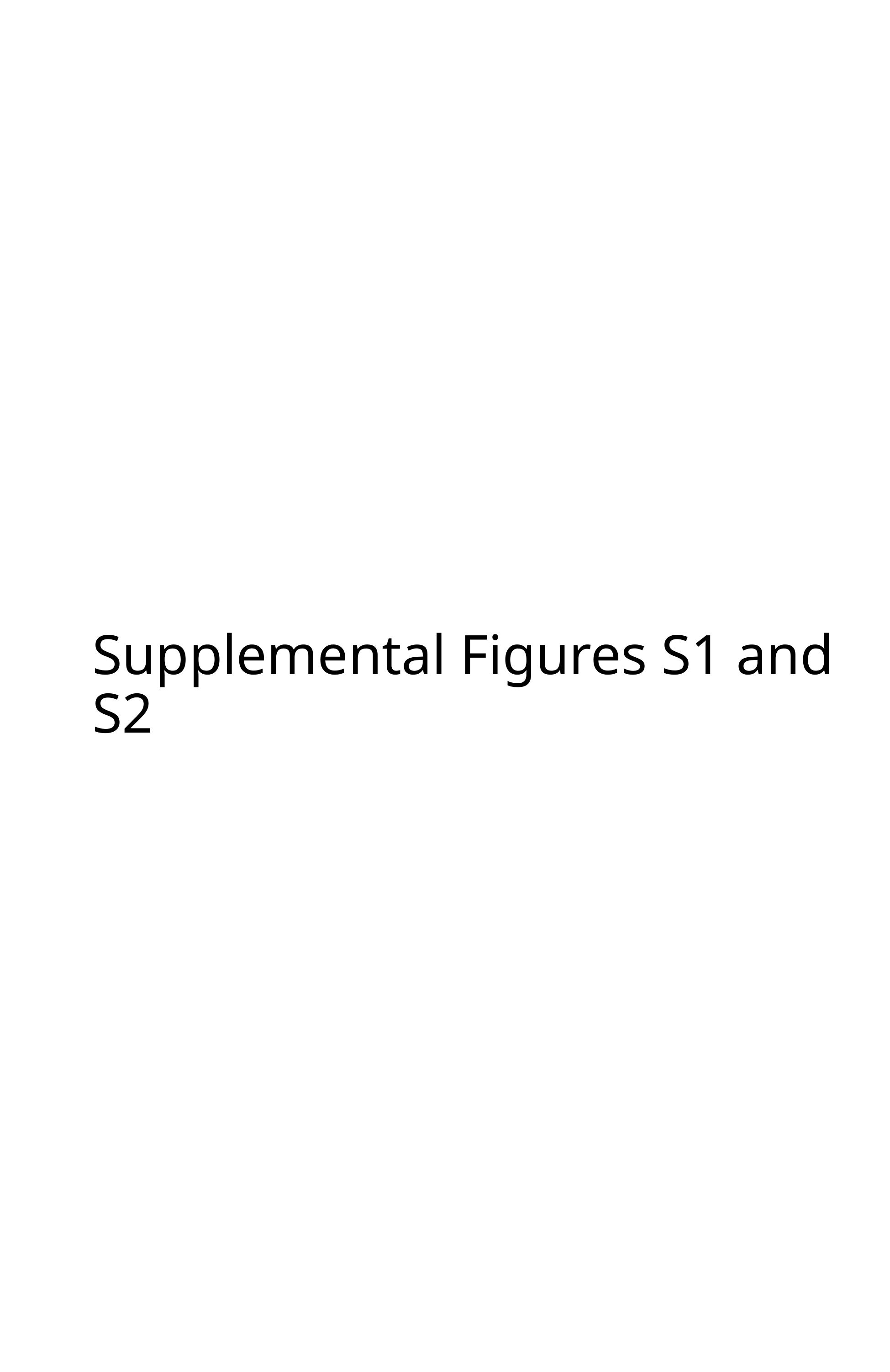

#### Supplemental Figures S1 and S2

### Slide 2
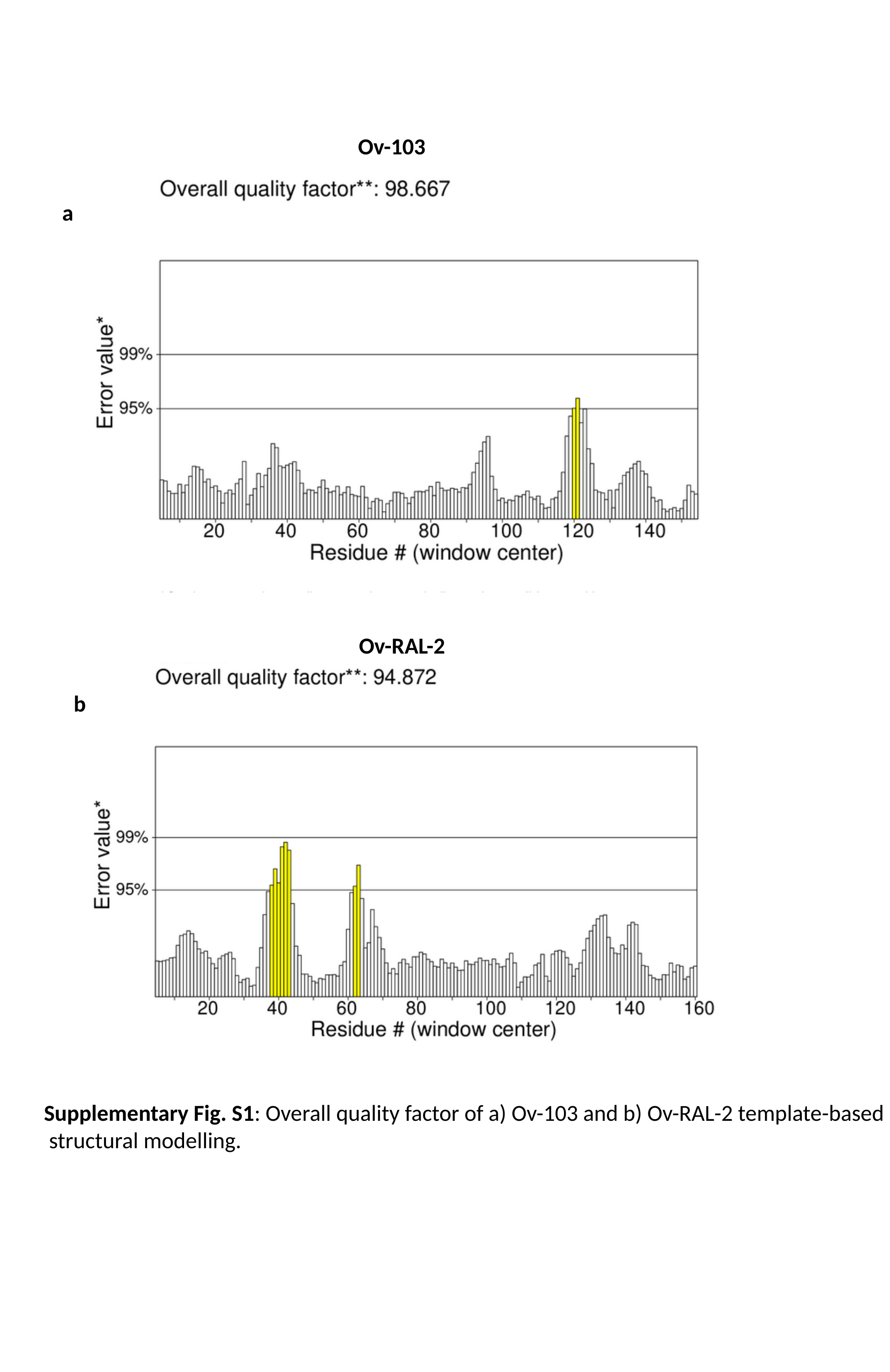

Ov-103
a
Ov-RAL-2
b
Supplementary Fig. S1: Overall quality factor of a) Ov-103 and b) Ov-RAL-2 template-based
 structural modelling.

### Slide 3
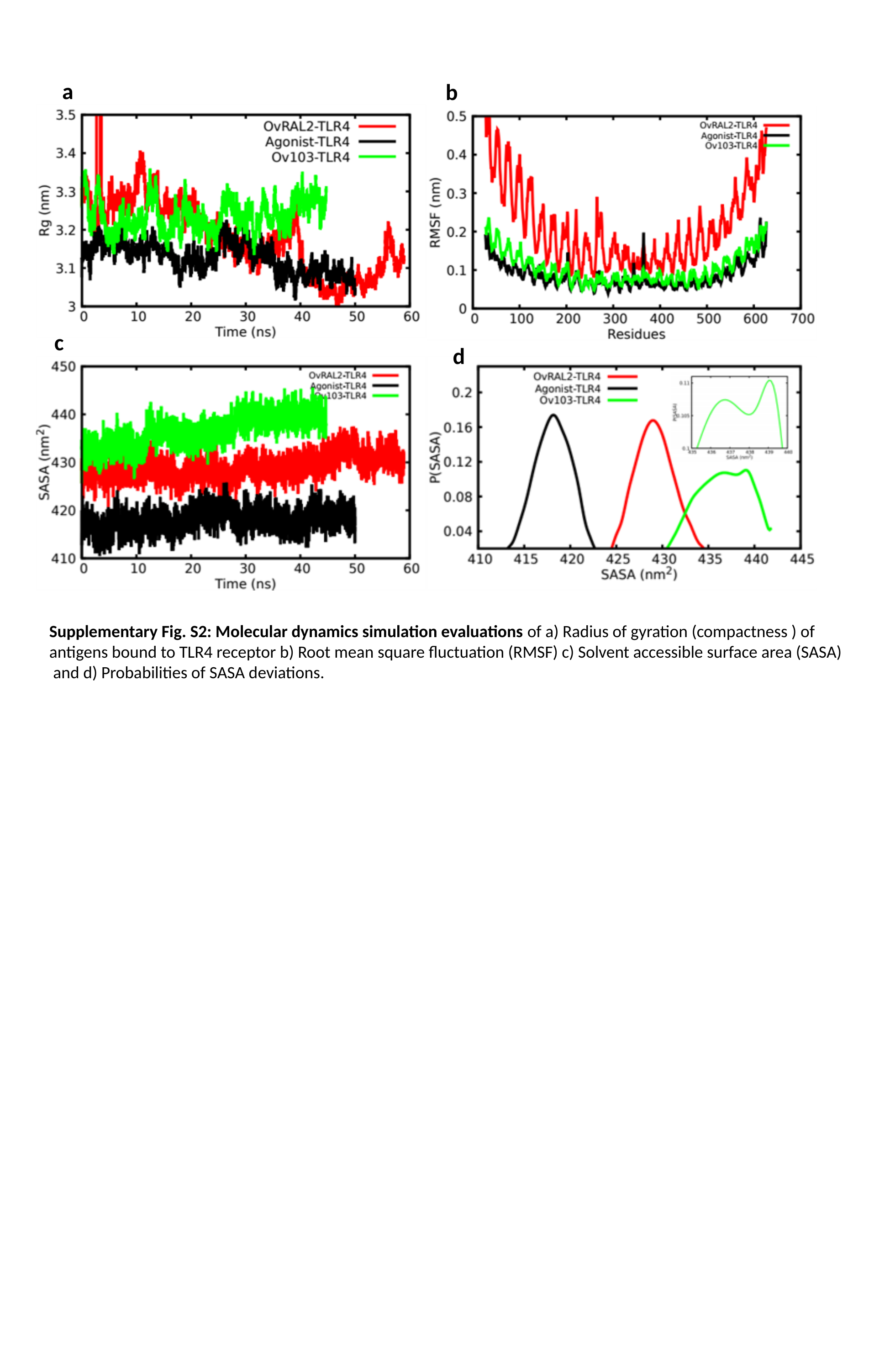

a
b
c
d
Supplementary Fig. S2: Molecular dynamics simulation evaluations of a) Radius of gyration (compactness ) of
antigens bound to TLR4 receptor b) Root mean square fluctuation (RMSF) c) Solvent accessible surface area (SASA)
 and d) Probabilities of SASA deviations.
